# Cell-extracellular matrix interactions in the fluidic phase direct the topology and polarity of self-organized epithelial structures

**DOI:** 10.1101/2020.05.12.090068

**Authors:** Mingxing Ouyang, Jiun-Yann Yu, Yenyu Chen, Linhong Deng, Chin-Lin Guo

## Abstract

*In vivo,* cells are surrounded by extracellular matrix (ECM). To build organs from single cells, it is generally believed that ECM serves as scaffolds to coordinate cell positioning and differentiation. Nevertheless, how cells utilize cell-ECM interactions for the spatiotemporal coordination to different ECM at the tissue scale is not fully understood. Here, using *in vitro* assay with engineered MDCK cells expressing H2B-mCherry (nucleus) and gp135/Podocalyxin-GFP (apical marker), we show in multi-dimensions that such coordination for epithelial morphogenesis can be determined by cell-soluble ECM interaction in the fluidic phase. The coordination depends on the native topology of ECM components such as sheet-like basement membrane (BM) and type I collagen (COL) fibers: scaffold formed by BM (COL) facilitates a close-ended (open-ended) coordination that leads to the formation of lobular (tubular) epithelium. Further, cells form apicobasal polarity throughout the entire lobule/tubule without a complete coverage of ECM at the basal side, and time-lapse two-photon scanning imaging reveals the polarization occurring early and maintained through the lobular expansion. During polarization, gp135-GFP was converged to the apical surface collectively in the lobular/tubular structures, suggesting possible intercellular communications. Under suspension culture, the polarization was impaired with multi-lumen formation in the tubules, implying the importance of ECM biomechanical microenvironment. Our results suggest a biophysical mechanism for cells to form polarity and coordinate positioning at tissue scale, and in engineering epithelium through cell-soluble ECM interaction and self-assembly.

## Introduction

One feature in epithelial development and regeneration is the spatiotemporal coordination of cell positioning and differentiation(Quintin et al., 2008). In these processes, cells develop apicobasal polarity and form lobular or tubular sheets(Bryant and Mostov, 2008; Martin-Belmonte and Mostov, 2008). Most remarkably, they can coordinate the orientation of polarity throughout the entire tissue(Martin-Belmonte and Mostov, 2008). Loss of such coordination is often a hallmark of tumors(St Johnston and Ahringer, 2010). As such, understanding how epithelial cells coordinate their positioning and polarization is not only essential for developmental biology and regenerative medicine, but also important for cancer biology. The transmembrane glycoprotein gp135 (also called Podocalyxin) is often used as an apical marker located at the luminal surfaces(Bryant et al., 2010; Cait et al., 2019), and Rab small GTPases mediate the direct transportation of gp135 and apicobasal polarization(Klinkert et al., 2016; Mrozowska and Fukuda, 2016). Over the past few decades, studies on epithelial morphogenesis have indicated the importance of cell-cell adhesions and cell-extracellular matrix (ECM) interactions(Blaschke et al., 1994; Bryant et al., 2010; Dickinson et al., 2011; Mailleux et al., 2008; Martin-Belmonte et al., 2007; Nelson, 2003; O’Brien et al., 2001; Rozario and DeSimone, 2010; Yu et al., 2005). The differentiation of epithelial cells in which non-polarized cells form polarized epithelium depends on ECM components(Blaschke et al., 1994). It was shown that breast epithelial cells differentiate into tubules in type I collagen (COL)(Dhimolea et al., 2010; Wozniak et al., 2003), while they form lobular acini in basement membrane (BM, mimicked by Matrigel in experiments)(Muthuswamy et al., 2001; Paszek et al., 2005). Our previous study further showed that cell-collagen interaction permits a long-range morphogenetic coordination at sub-millimeter scale(Guo et al., 2012). Nevertheless, how epithelial cells utilize cell-ECM interaction to coordinate their positioning and polarization in response to different ECM components at the whole-tissue scale is not fully understood.

To form long-range coordination, it is generally believed that ECM can serve as scaffolds to guide cell positioning and polarization(Rozario and DeSimone, 2010). Cells constantly secrete soluble ECM molecules and degrade existing ECM scaffolds into soluble fragments *in vivo.* Theoretically, these soluble forms of ECM can be assembled into scaffolds through two processes. The first is that they self-assemble into new scaffolds or re-incorporate into pre-existing scaffolds. Alternatively, soluble ECM can interact with cells which serve as nucleation cores to assemble ECM scaffolds. An example is the development of renal tubules where BM components are found to be dynamically assembled around the pre-tubular structure(Ekblom et al., 1980). Apicobasal polarization is a typical process during epithelial tubulogenesis, and consistently with the role from ECM, the cellular mechanism involving integrin and RhoA signaling pathways has been identified in triggering the polarity formation(Bryant et al., 2014; Kim et al., 2015).

Here, we study whether ECM scaffolds created by ECM self-assembled hydrogel or by cell-mediated assembly play the primary roles in the formation and coordination of epithelial morphogenesis. These two processes can hardly be decoupled *in vivo* or through the conventional ECM reconstitution approaches. Our recent work demonstrated that cell motion promotes fibrillary assembly of soluble COL, in supporting the role of cells in ECM scaffold generation(Wang et al., 2020). We therefore use the *in vitro* open-system assay, and found that spatiotemporal coordination in epithelial morphogenesis and polarization can occur on cell-assembled ECM in the fluidic phase rather than pre-assembled ECM in the solid phase. The coordination depends on native topology of the ECM components such as basement membrane (BM) and type I collagen (COL). Further discovered during tubulogenesis, the apicobasal polarization proceeds in a collective way along the axis of the tubule, implying intercellular communications within the cell groups. Our results suggest a potential mechanism which cells can use to form polarity and coordinate morphogenesis *in vivo*, and a strategy to engineer epithelial structures through self-assembly *in vitro*.

## Results

### Cell-ECM interactions in the fluidic phase for polarized epithelium formation

We first examined if cells can develop coordinated polarity on ECM scaffolds formed by ECM self-assembly without any soluble ECM. Madin-Darby Canine Kidney (MDCK) cells were used in this study, which are a popular model cell line for epithelial morphogenesis and apicobasal polarity formation regulated by Rab GTPases(Martin-Belmonte and Mostov, 2008; Mrozowska and Fukuda, 2016). To track the development of apicobasal polarity, we engineered MDCK cells stably expressing mCherry-conjugated histone H2B (H2B-mCherry) and GFP-conjugated gp135 (gp135-GFP). Here, H2B-mCherry is used to indicate cell nucleus(Sato et al., 2010), whereas the apical marker gp135 is used to indicate cell polarization(Bryant et al., 2010; Meder et al., 2005). To define polarity, we noted that in polarized epithelium, cells are organized into a surrounding and continuous monolayer structure with gp135 primarily confined at their apical surfaces(Bryant et al., 2010), which are located at the inner space of the organization (illustrated in Fig. 1a with the experimental data shown in Fig. 1b). Thus, we used the spatial distribution of H2B-mCherry with respect to gp135-GFP to define and track the formation of polarity. The polarization process here refers to the convergence of diffusive gp135-GFP to the apical surface of the lumen or to the intercellular region between cell nuclei. To initiate cell density-dependent long-range coordination, we adopted the cell density characterized and optimized in our previous study (Guo et al., 2012).

**Figure 1.**
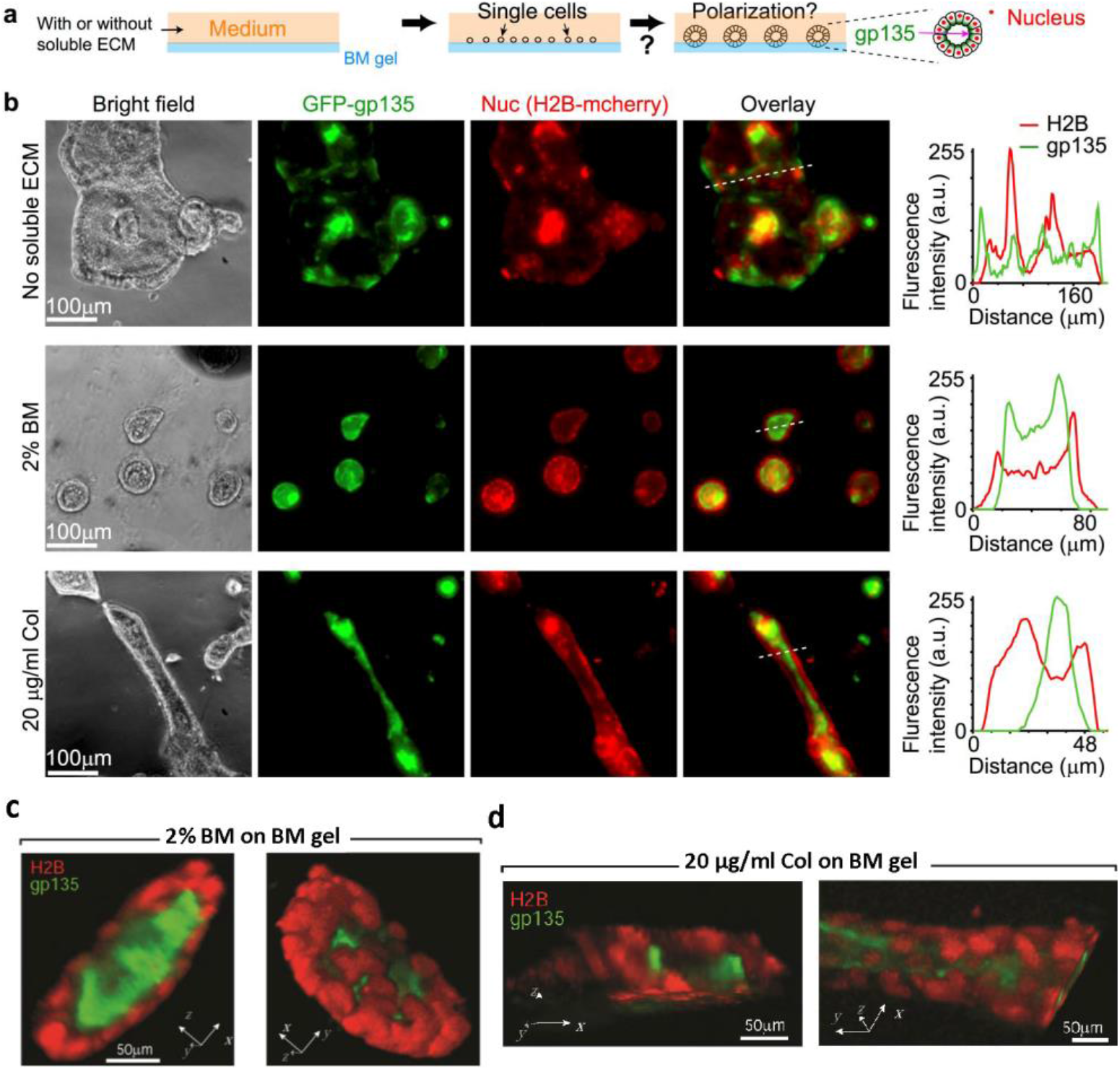
Soluble ECM components are required to form epithelial polarization on Matrigel. **(a)** Experimental setup. MDCK cells were cultured on the top of basement membrane (BM) gels with or without soluble ECM components in the medium. Cells express H2B-mCherry and gp135-GFP to indicate their nucleus and polarity, respectively. Note the distribution of nucleus with respect to gp135 in polarized lumens (on the right). **(b)** Left in each row: Represented phase contrast (bright field) and 3-D projected fluorescent images (gp135: green, H2B: red, and their overlay) for cells seeded on BM gels and cultured for 12 days (seen in Methods for 3-D projected imaging and processing). The medium contained (top row) no soluble ECM, (middle row) 2% BM, or (bottom row) 20 μg/ml type I collagen (COL). Nuc: cell nucleus. Right in each row: Fluorescence intensities of H2B-mCherry and gp135-GFP along the indicated white dotted lines (from left to right). The overlay of the red and green curves indicates the positions of cell nucleus (red) with respect to the apical marker (green). a. u.: arbitrary unit. **(c & d)** 3-D view of the polarized lobular and tubular structures. The images from confocal scanning microscopy (with 20x objective) were reconstructed into 3-D structures. The 3-D views showed the relative position of cell nucleus (red) and apical marker gp135 (green) in the closed lobular (c) and elongated tubular (d) structures.

After long-term culture, epithelial cells including MDCK cells can secrete ECM molecules(Ecay and Valentich, 1992; Parry et al., 1985; Yu et al., 2005). We therefore used open systems to dilute and/or remove secreted ECM by changing medium every day or every other day. Cells were cultured on pre-assembled BD Matrigel gels (to mimicking basement membrane (BM), the primary matrix components underlying polarized epithelium *in vivo*) open to a large medium space that contained no soluble ECM (Fig. 1a). Under this condition, cells grew and merged into big clusters (hundreds of micrometers in diameter) without forming general apicobasal polarity (Fig. 1b, top row). In order to visualize the cultured epithelium, 3-D projection of the epifluorescence images taken at 21 planes with a step size of 4.5 μm was obtained by collecting pixels with the maximal intensity through the entire z-stack into a single plane, which is further explained in the Methods.

We then examined if cell-ECM interactions in the fluidic phase is required for polarity formation. Two types of medium were prepared: medium with BM (2%, diluted Matrigel solution from ~10 mg/ml stock concentration), and medium with COL (20 μg/ml). From the manufacturer, the major components of Matrigel are laminin (~61%), collagen IV (~30%) and Entactin (~7%). The BM concentration was adapted from studies by our group and others (Blaschke et al., 1994; Bryant et al., 2014; Guo et al., 2012; Klinkert et al., 2016; Muthuswamy et al., 2001), and the COL concentration at 20 μg/ml was designed to match the similar mass concentration of 2% BM. Here, we used low concentrations of soluble ECM components to prohibit their fast self-assembly into hydrogel-like. This allowed cells to live in a fluidic or semi-fluidic condition and interact with soluble ECM during their proliferation. Here semi-fluidic condition referred to cells/clusters culturing on solid Matrigel scaffold with direct exposure to the medium. In response to soluble BM, cells were found to form spherical, lobular structures with coordinated polarity (Fig. 1b, middle row), whereas for soluble COL, they formed polarized, tubular structures (Fig. 1b, bottom row). In both cases, gp135 was confined within the apical area (Fig. 1b, middle and bottom rows, right). 3-D view of these cultured lobular and tubular structures from confocal scanning microscopy further confirmed that cells grew into fine 3-D structures with gp135 confined at the apical surfaces (Fig. 1c&d).

### Dynamics of cell positioning and polarization dependent on cell-ECM interactions

Having shown that cell-BM (COL) interactions in the fluidic phase leads to the formation of polarized lobules (tubules), we examined how cells coordinate their positioning and polarization in response to soluble BM (COL). Same as the setups in Fig. 1a, cells were seeded on 3-D BM with or without soluble ECM in the medium, and the dynamic processes of cell aggregation and polarization were continuously recorded by time-lapse imaging in the following days.

We first examined cellular dynamics on BM gels in the presence or absence of soluble BM. Without soluble BM, time-lapse microscopy revealed that individual cells proliferated into small clusters, which continuously grew and merged with each other without general polarity formation (through the 5-day period of observation, Movie 1) (Fig. 2a, Movie 1). By contrast, with soluble BM, most clusters stopped merging and became polarized on the 2nd-3rd day (Fig. 2b). Here, the timing of polarization was defined by the conversion of gp135 from the outer layers to the inner areas of clusters/cysts (Fig. 2b, and Movie 2). The lumen cultured by MDCK cells (H2B-mCherry/GFP-β-actin) displays actin ring at the apical side (Fig. 2c), which was consistent with the previous report(Ivanov et al., 2008).

**Figure 2.**
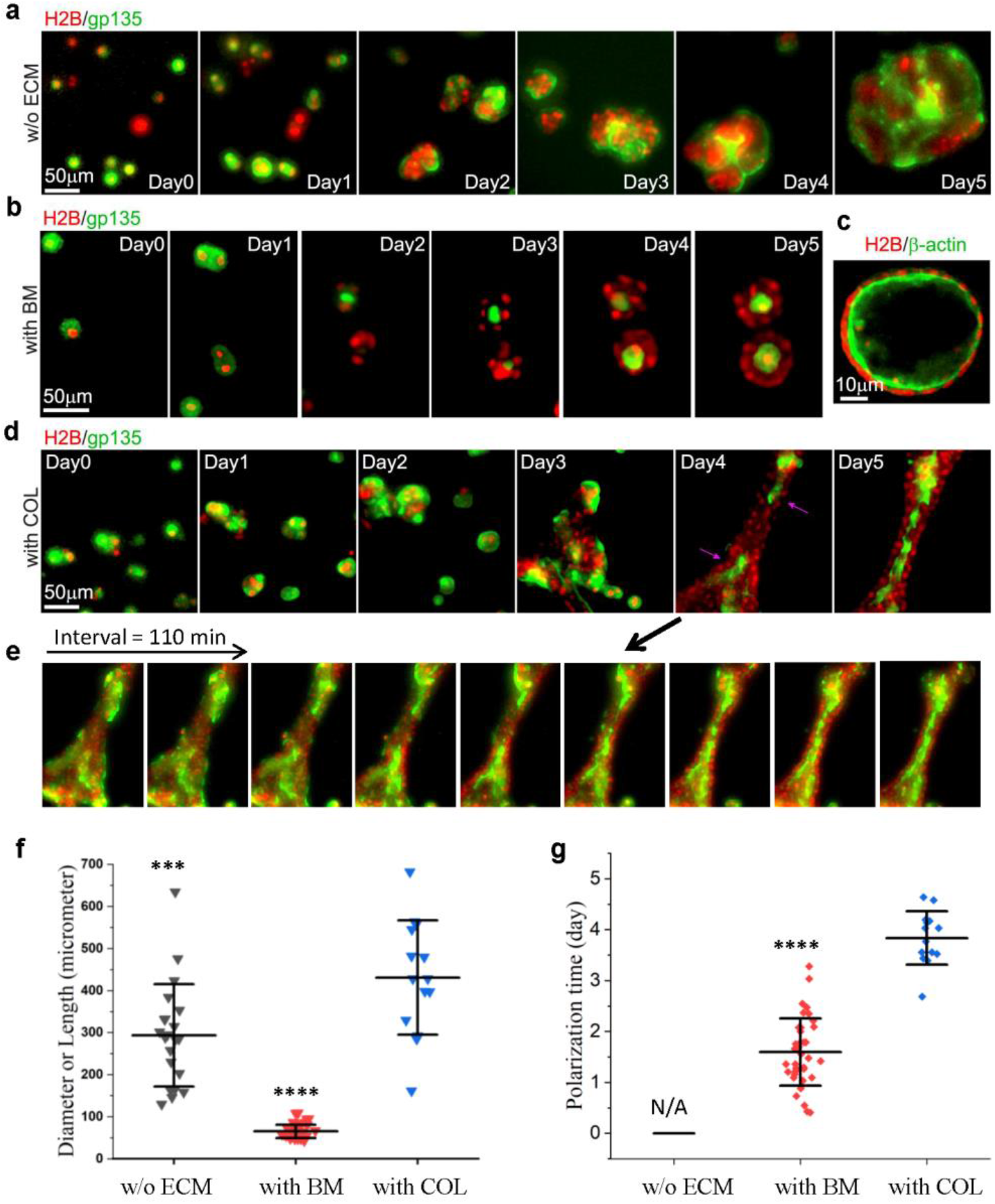
Distinct dynamics of cell coordination in response to soluble BM/COL. MDCK cells express H2B-mCherry and GFP-gp135 or β-actin-GFP to indicate cell nucleus and apicobasal polarity, respectively. Cells were cultured on BM gel with or without soluble ECM in the medium and positioned on the microscope by change with fresh medium every day. Time-lapse fluorescence images were taken with 22 min interval time per cycle in the following days. **(a & b)** Represented time series of 3-D projected epifluorescence images (gp135: green, H2B: red) for cells in medium (a) without ECM (n >10), (b) with 2% basement membrane (BM) components (n >10) (also seen in Movies 1&2, interval = 22 min). **(c)** Represented optical sectioning from scanning microscopy (β-actin: green, H2B: red) to demonstrate lumen formation with polarized actin distribution after culturing cells in medium containing 2% BM for 14 days. **(d)** Represented time series of 3-D projected epifluorescence images for cells in medium with 20 μg/ml COL (also seen in Movie 3) (n=14). The purple arrows indicate the polarization initiating from the local regions of the tubule. **(e)** The collective apicobasal polarization along the tubular axis (n=6). The time-sequence images (interval: 22min*5 = 110min) showed the spatial distribution of gp135 relative to cell nuclei (H2B) during the polarization. (**f**) Size quantification of the diameters (along the long axis) of unpolarized clusters without ECM (n=21) and polarized lobular lumens with BM (n=50), and the length of polarized tubes with COL (n=14) around Day 5. (**g**) The timing quantification when lumens (n=38) or tubules (n=14) got polarized under culture with 2% BM or 20 μg/ml COL in the medium. N/A refers to “not applicable for polarization” without ECM in the medium. The data quantification (mean±S.D.) was performed by using ImageJ and Origin. *** and **** represent significant difference with P < 10^-2^ and 10^-6^ in comparison to the group “with COL” from Student’s t-test analysis.

We next examined the cellular dynamics on BM gels with soluble COL. Similar to the observation with soluble BM (Fig. 2b), cells were found to form clusters which proliferated and coalesced. However, the coalescence did not stop on the 3^rd^ day but instead clusters continued to fuse into a long-range (>200 μm), tubule-like structure (Fig. 2d). By then, cells started to polarize through a collective conversion of gp135 from the outer layer to the inner area of tubule (Fig. 2d, Movie 3). From the time-sequence images, the polarization in the presence of soluble COL may proceed collectively along the axis of tubule in the same cluster in which the spatial localization of gp135 turned over from the boundary to the center of the tubule along the polarization process (Fig. 2e, Movie 3). The way of the tubular assembly had impact on whether there was typical collective polarization (~30%). During the coalescence, we often observed mutual attraction of clusters on the field of views under the time-lapse microscopy (Movie 3). Such prolonged coalescence was not observed in the case where cells were surrounded with soluble BM, suggesting that it is a feature of COL-cell interactions.

From the size quantification, unpolarized clusters (w/o ECM) grew into hundreds of micrometers in diameter, and polarized lobular lumens (with BM) were generally within 100 μm, while the length of the polarized tubes (with COL) reached ~400 μm at average (Fig. 2f). The polarization time was much different between lobular lumens (generally within 2 days) and tubes (~4 days) (Fig. 2g), which suggests different morphogenetic mechanism coordinated with BM and COL.

### Cell polarization in the fluidic phase with ECM assembly

If the coordination of cell positioning and polarization requires the formation of ECM scaffolds, the results above suggest that it is the scaffold mediated by cell-ECM interaction in the fluidic phase that determines epithelial morphology and coordinates polarity. To see how such scaffolds are formed and affect cell positioning and polarization, we performed immuno-staining of laminin (a major component in BM(Kalluri, 2003)) and COL on polarized lobules and tubules.

We first examined how laminin is distributed on cells seeded on BM gels with or without soluble BM. In the presence of soluble BM, condensed laminin was found at the outer layers of clusters after 3 days of culture (Fig. 3a), and the density of laminin increased during the culture time (Fig. 3b), indicating the laminin assembly along with the cluster growth. After 7 days, most clusters were found to form polarized lobules surrounded by dense laminin (Fig. 3a). By contrast, laminin was not found at the outer layers of clusters in the absence of soluble BM (Fig. S1), where clusters continuously grew and coalesced without polarity formation (Movie 1), suggesting that laminin assembled around clusters might block coalescence and induce polarization.

**Figure 3.**
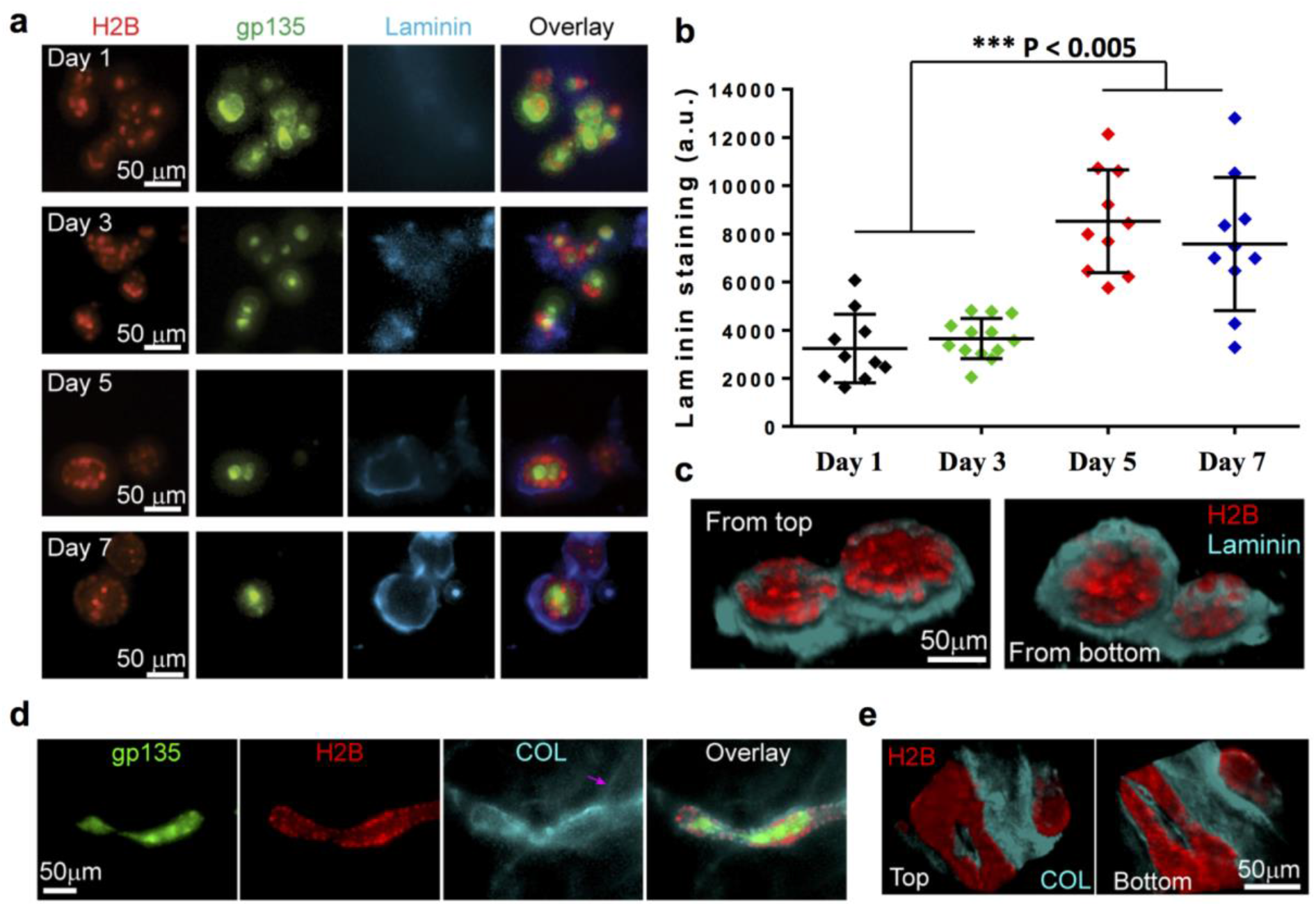
Cell polarization in the fluidic phase with and without direct cell-ECM contact. **(a, b)** Lateral assembly of laminin around the luminal structure. MDCK cells were cultured for 1-7 days on BM gels with 2% BM in the medium, followed by laminin immuno-staining. (a) Represented fluorescent images of gp135-GFP (green), H2B-mCherry (red), laminin staining (cyan), and their overlay. (b) Fluorescence quantification of laminin staining on the cultured MDCK clusters (a). The graph data with individual dots (mean±S.D.) show the average fluorescence intensity over the entire clusters with typical staining signals. a.u.: arbitrary unit. **(c)** Represented images of 3-D reconstructed, top and bottom views of cell nucleus (red) and laminin (cyan) on the spherical lobules after 12-day culture (also seen in Movie 4). **(d)** Represented fluorescent images of gp135-GFP (green), H2B-mCherry (red), COL staining (cyan), and their overlay of the tubules (n=15). MDCK cells were cultured for 12 days on BM gels with COL (20 μg/ml) in the medium, followed by COL immuno-staining. Note the formation of linear structure of COL (pink arrow). **(e)** Represented images of 3-D reconstructed, top and bottom views of cell nucleus (red) and COL (cyan) on the tubule after 12-day culture. For (c & e), images were taken by confocal scanning microscope and created by ImageJ 3-D reconstructions. Note the lateral condensation of laminin (COL) around the lobules (tubule) (outlined by nucleus) and the absence of laminin (COL) assembly on the top and bottom.

In the conventional model, the development of epithelial apicobasal polarity requires coordinated cell-ECM interaction at the basal side (Mostov et al., 2005; O’Brien et al., 2001) and cell-cell interactions (Bryant and Mostov, 2008; Dickinson et al., 2011) at the lateral side of each individual cell. To ascertain if this is the case in our in vitro assays where cell-ECM interactions at the fluid phase appeared to play the deterministic role in apicobasal polarity formation, we examined the distribution of laminin around the clusters undergoing polarization. Cells on BM gels were cultured for 12 days with soluble BM to form polarized lobular lumens. Confocal scanning microscopy was then performed to construct 3-D views of laminin and lumens with the lumens outlined by H2B-mCherry signal from cell nucleus. Surprisingly, no complete, uniform assembly of laminin around the lobular lumens was found. Instead, assembled laminin was found to form a lateral platform at the lateral side of each lumen, rather than on the top (facing the medium) or bottom (facing the BM gel) (Fig. 3c, and Movie 4), whereas cells at the entire lumen were polarized (Fig. 1c). These results might also suggest that cells at the top of lumen could maintain polarity in the absence of direct cell-ECM contact. The laminin platform appears as a 2-D network compatible with the native topology of BM(Ingber, 2006).

To see if cells can also polarize without direct cell-ECM contact in the presence of soluble COL, we examined the distribution of COL around the clusters that underwent polarization on BM gels with soluble COL in the medium. After 12 days of culture, cells formed tubules and found associated with linear structures of COL (Fig. 3d, pink arrow). Confocal scanning microscopy revealed that COL was condensed at the lateral sides, but not the top or bottom of tubules (Fig. 3e), and similar to the results in the case with soluble BM (Fig. 3c), cells at the entire tubules were polarized (Fig. 3d). These suggest that those cells without ECM interaction might maintain their polarity through cell-cell interactions in the lobules and tubes.

### The expansion of dividing cells into fine spherical lobular structure under polarization

To further understand how cells maintained apicobasal polarity at the entire epithelial structures with only partial cell-ECM interactions during the culture, we carried time-lapse confocal imaging to observe the growth of dividing cells into polarized lobules. This experiment had certain technical challenges: due to the smaller field by confocal imaging than the wide-field epi-microscopy, the motile cell samples could move out of the views more likely; second, confocal imaging had higher photo-toxicity during the point-by-point scanning and z-stack imaging, which could cause damage on cell samples. To manage to get some time-lapse samples successfully, we tried to lower the excitation laser power (25 mW) and used long interval time (3 or 4 hours).

As the cell cluster growth shown in Fig. 4 with more detailed views in Movie 5, the cell sample started at two-cell stage, and continued to divide into 3 and 4-cell cluster without polarization (located on one plane) on the next day (Fig. 4a); late on the third day, the polarization was occurring in one cluster (not sure whether the same one from the beginning), and in the following days, the cluster continued to grow and expand into fine 3-D lobular structure with maintained polarization, suggesting the possibility of cell divisions under polarized status in the lobule (Fig. 4b). It is noted that the polarization of the cell cluster may occur in a collective way and gp135 was turned over into the cluster from one local side (Fig. 4b, the first panel), which also indicates possible intercellular communications. These time-sequence images indicate that the cell cluster got polarized at early stage, and grew into fine-constructed 3-D lobule under maintained polarization. The result may help explain how cell polarization occurred on the entire lobules although there was only partial laminin coverage on the basal side (Fig. 3c). Similar mechanism may be extended to the observed tubular structure with partial COL coverage. In considering that this is a descriptive data with limited work, we didn’t have explanation why ECM components weren’t assembled on the top of the lobule, or dig out more insights at current stage.

**Figure 4.**
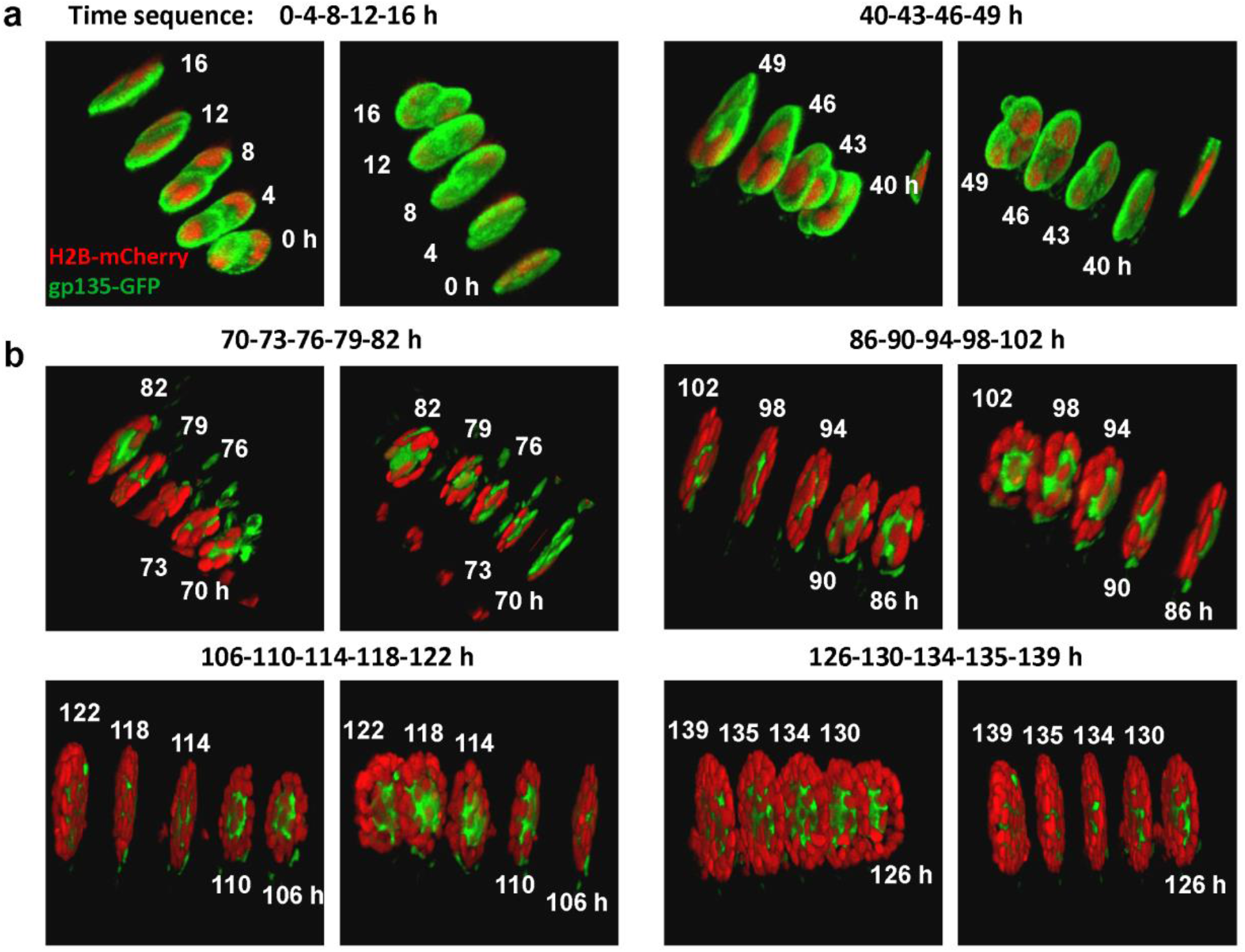
The expansion of dividing cells into polarized lobule. The MCDK cells expressing gp135-GFP and H2B-mCherry were seeded on BM gel with 2% BM in medium, and two-photon confocal imaging started one day later. The wavelengths of excitation light were 890 nm for GFP and ~1100 nm for mCherry, the step size in z-direction was 1.0 μm. Time-lapse imaging was taken with the interval time of 3 or 4 h. Medium was changed every day along with focus/position corrections. Acquired confocal images were processed by two channels (GFP, mCherry) overlay and 3-D reconstructions. Each image here shows the 3-D view of 4 or 5 time-points at the labeled time (in hours, and the starting time was set as zero), and two images for each time point were displayed from different angles (generally from the top and the bottom views). **(a)** The polarity distribution at early stage in the cluster with a few cells. **(b)** The polarity distribution during growth of the cluster into fine 3-D lobular structure. More detailed 3-D views are shown in Movie 5. A note: the images of (a, b) were acquired at the same position in one experiment, but may not be from the exact same sample during the re-focusing process as cells were motile at the early stage.

### Culture of lobular and tubular epithelial structures under suspension conditions

The results above suggest two distinct, ECM-dependent processes to coordinate epithelial morphogenesis in the fluidic phase. The first is that cells recruit soluble BM components which are known to form branched networks(Ingber, 2006), to create a closed-end (i.e., restricted) scaffold surrounding individual cluster. Such scaffold provides a physical barrier to block the coalescence of clusters and allow them to proliferate and polarize within the restricted space (Fig. 3a). The second process is that cells recruit soluble COL, which is known to form linear, bundled fibers(Kadler et al., 2008; Starborg et al., 2008), to create an open-ended (i.e., unrestricted) scaffold (Fig. 3d). In contrast to the first one, scaffold formed by soluble COL allows clusters to continuously merge with one another through long-range interactions in the fluidic/semi-fluidic phase.

These two processes were observed in the assays where cells were supported by pre-assembled ECM, i.e., a solid phase. To ascertain whether the solid phase is absolutely required in soluble ECM-mediated epithelial cell polarization and morphogenesis, we repeated the assay in a suspension system where agarose gel was used to replace the solid phase and minimize cell-substrate interaction (Fig. 5a) (details seen in Methods). Cells (final concentration of ~1×10^4^ cells/ml) were mixed with medium containing no ECM (as the control) or soluble ECM components (2% BM or 20 μg/ml COL), spread in the suspension system, and examined for polarization after 7-14 days.

**Figure 5.**
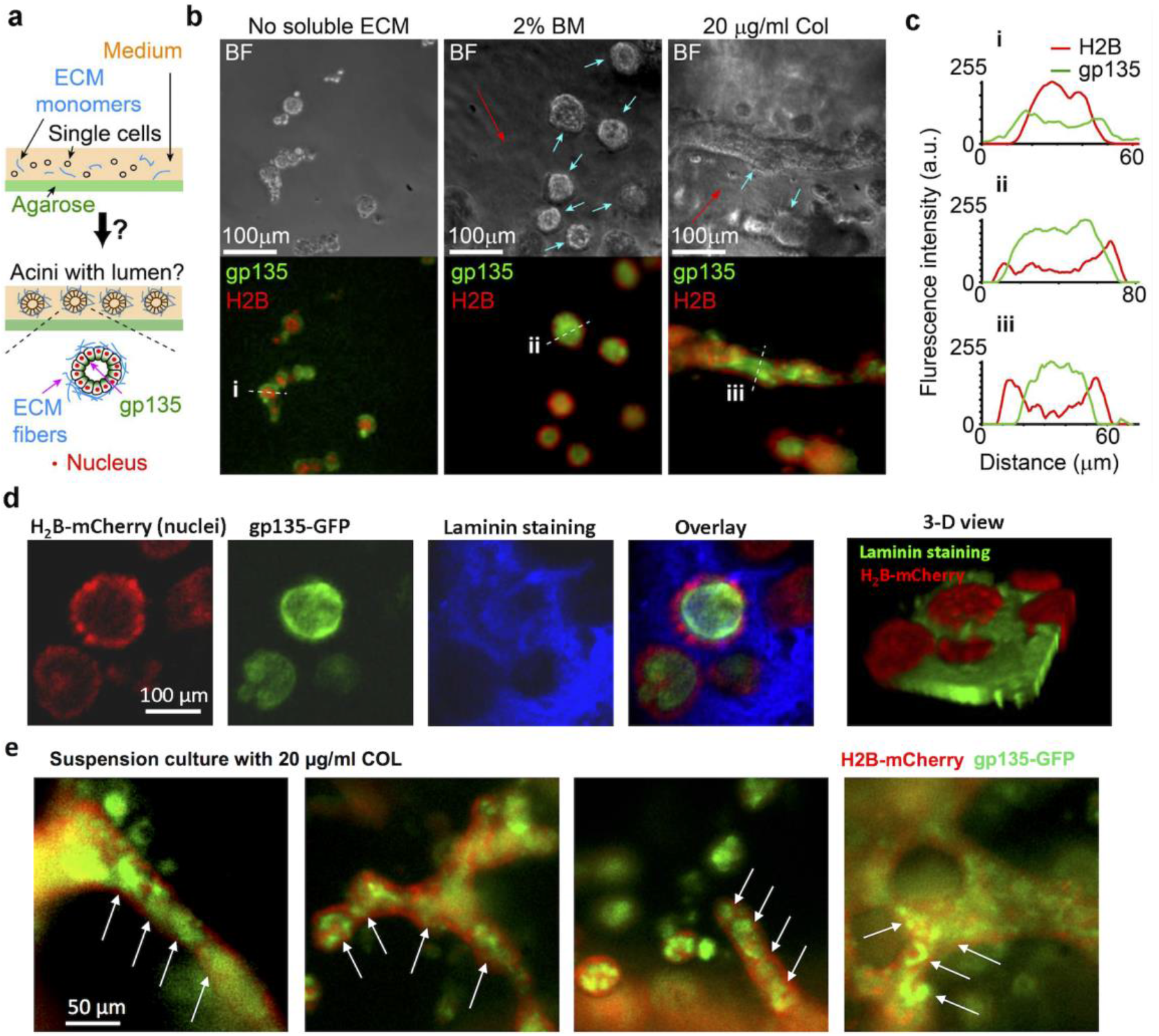
Cells form tubular and lobular structures above non-adhesive substrates. **(a)** The design of suspension culture on agarose gel. Details are described in Methods. **(b)** Representative bright-field and 3-D projected epifluorescence images (Green: gp135, Red: H2B) for MDCK cells cultured in medium containing no ECM materials (Left), 2% BM (Middle), or 20 μg/ml COL components (Right) above 1% agarose gel for 12 days. Initial cell density: ~10^4^ cells/ml. To enhance visualization, out-of-focus signals were removed and only fluorescence signals from cells near the focal plane (indicated by blue arrows) were shown. The red arrows indicated the scaffolds self-assembled from soluble BM or COL. **(c)** Fluorescence intensities of H2B-mCherry and gp135-GFP along the white dotted lines (from left to right) in the indicated H2B/gp135 overlay. The cell nucleus (red) located outside the apical marker (green) indicates polarity formation. **(d)** The 3-D projected images of spherical lobules from suspension culture with 2% BM (~12 days). The laminin staining (blue) indicated the assembled scaffolds from soluble BM under suspension. The images were taken by confocal microscopy (20x objective). 3-D projection of the images was obtained by collecting pixels with the maximal intensity through the entire z-stack into a single plane, and the 3-D view was constructed by the software ImageJ. **(e)** Formation of multi-lumen polarity on the tubules under suspension culture. MDCK cells were cultured above agarose gel with 20 μg/ml COL in the medium for 12 days. The arrows indicate some of these small lumens on the tubules.

Without soluble ECM components, cells in the suspension system were found to form clusters with a reverse polarity (i.e., gp135 located at the outer boundaries of clusters, Fig. 5b left panel), which is consistent with the previous finding (Wang et al., 1990). By contrast, cells mixed with soluble ECM components formed structures of polarized epithelium (Fig. 5b, middle and right). Specifically, cells formed polarized lobular structures with BM (Fig. 5b middle panel) and polarized tubular structures with COL (Fig. 5b right panel), with the epithelial polarity indicated by the spatial distributions of H2B-mCherry and gp135-GFP signals (Fig. 5c(i-iii)). We noticed that under suspension culture with COL, separated small rooms (multi-lumens) were formed in the tube during polarization (Fig. 5b&e), instead of homogenous polarity through the tube on solid Matrigel (Fig. 1b), which may reflect certain different ECM biomechanical microenvironments in the two culture systems. Specifically, the cells & ECM scaffolds floated in the medium during suspension culture, and the ECM scaffolds were also soft enough for antibody staining, whereas the ECM scaffold was bounded to glass surface during culture on the solid Matrigel which is also resistant to antibody staining.

Laminin immuno-staining showed the distribution of ECM scaffold around the polarized lumen (Fig. 5d), and laminin only covered the lateral sides of lobules, not at their top or bottom sides (Fig. 5d (right)), which seemed similar to the results found in the lobules cultured on top of BM gels (Fig. 3c). It was noted that soluble BM and COL in the medium could form scaffolds alone (Fig. 5b&d), which assisted the formation of polarized epithelium. Thus, cell-ECM interactions are required to initiate epithelial morphogenesis and create polarization at the tissue scale.

## Discussion

Emerging evidence suggests that cell-ECM interactions are crucial in regulating tissue development, homeostasis, and repair(Behonick and Werb, 2003; Klinkert et al., 2016; Sequeira et al., 2010). Such interactions can occur on the solid phase (i.e., the existing ECM scaffold) and in the fluidic phase where cells interact with soluble ECM components. Here, we showed the following findings based on the engineered MDCK cells stably co-expressing H2B-mCherry (nucleus) and gp135-EGFP (apical marker): (1) the topology of native ECM components from cells-mediated assembly helps define the morphology of epithelium tissue; (2) during tubulogenesis in the presence of soluble COL, the apicobasal polarization proceeds in a collective way along the axis of the tubule; (3) in the fluidic phase, cells can form apicobasal polarity throughout the entire lobule/tubule with ECM only partially covering the basal surfaces (i.e. without ECM assembly at the top/bottom sides), and polarization occurred at early stage and was maintained through the entire lobular expansion; (4) under suspension culture with COL, the polarization was impaired by forming multi-lumens on the tubules, suggesting the importance of ECM biomechanical microenvironment for tubulogenesis.

### Self-organization of polarized epithelium in the fluidic phase

First in our culture system, it is the scaffold formed by cell-ECM interactions in the fluidic phase plays important roles in epithelial polarization and morphogenesis. This enables the self-assembly of polarized epithelium. We note that 3-D culture embedded in BM gels can form polarized lobules(Blaschke et al., 1994), and has been used to study the polarization of cells obtained from tubular organs including the kidney, the liver, and the prostate(Martin-Belmonte and Mostov, 2008; Tanimizu et al., 2007; Webber et al., 1997). Such 3-D culture is different from our open-system setup. In our setup, soluble ECM components secreted by cells are diluted, whereas in the 3-D culture, they remain trapped within the space occupied by the cells. In turn, cell-soluble ECM interactions are allowed in the 3-D culture. Indeed, ECM components secreted from cells or degraded from gels is required for epithelial polarization in the 3-D culture(Davis et al., 2007; O’Brien et al., 2001). Consistently in the modified 3-D culture (developed by Brugge et al.(Muthuswamy et al., 2001)) where cells are cultured on BM gels (mimicked by Matrigel in experiments), soluble BM is required for cells to form polarized lobules(Bryant et al., 2010; Datta et al., 2011; Kim et al., 2009; Martin-Belmonte et al., 2007). These observations highlight the criticalness of cell-ECM interactions from the fluidic phase for epithelial polarization.

In general, soluble ECM can be assembled into scaffolds by ECM self-assembly and/or by cell-mediated nucleation process. The extent of cell-soluble ECM interactions in scaffold assembly depends on the cell density and the soluble ECM concentration. In our open-system setup, the concentration of soluble ECM is set low to eliminate or attenuate ECM self-assembly. By contrast, the 3-D model in previous studies(Blaschke et al., 1994) used high concentrations of soluble ECM components to induce ECM self-assembly.

It is likely that ECM scaffolds assembled by cell-soluble ECM interactions possess a more physiological structure. For example, BM gels assembled in the absence of cells were found more resistant to laminin staining than those assembled by cells, as indicated by the difference of laminin staining (Fig. 3a), which might result from difference of porosity. Indeed, previous reports showed that immune-staining of cells in BM or COL gels requires a pre-cleavage of gels by ECM proteinases(Kim et al., 2009; O’Brien et al., 2001).

### Coordination of cell dynamics dependent on soluble ECM components

Our second finding is the distinct coordination of cell positioning and polarization in response to soluble BM (diluted Matrigel solution) and COL. With soluble COL, cell polarization was found to arise after the formation of elongated structures. By contrast, with soluble BM, cell polarization can occur without fusion of individual clusters. In addition, while cell polarization in soluble BM occurs almost simultaneously within the same cluster, cell polarization in soluble COL appears to follow a nucleation process in the cluster (Fig. 2b&d, Movies 2&3). This collective polarization proceeded along the epithelial tubule (Fig. 2e) might suggest existing intercellular signal communications at the cell-population level during tubulogenesis. The selection of topology in the polarized epithelial clusters appears to correlate with the native topology of BM and COL (i.e., lobular or linear/tubular).

It has been documented that cell-ECM interactions through integrin signaling are crucial in regulating apicobasal polarity in epitheliums(Manninen, 2015). By application of functional-inhibitory antibodies, it was reported that integrin β1 is involved in the polarity regulation including the apical localization of gp135(Ojakian and Schwimmer, 1994). Further studies of the downstream mechanism showed that integrin-linking kinase (ILK) modulates laminin assembly and regulates integrin-microtubule network for directional delivery of apical factors(Akhtar and Streuli, 2013; Rudkouskaya et al., 2014). Multiple groups also revealed that small GTPases Rac and Rho signaling act downstream of integrins to mediate and maintain appropriate epithelial polarity(Bryant et al., 2014; Li and Pendergast, 2011; Yu et al., 2005; Yu et al., 2008). Our work here added one piece of information to the scenario that biophysical factors like cells-mediated assembly of ECM structure may play a role in directing the epithelial topology and polarity during morphogenesis. At this stage, nevertheless, we didn’t investigate how molecular signals from cell-soluble BM/COL interactions lead to distinct dynamics in polarity formation.

### Epithelial polarization with or without direct cell-ECM contact

Our third finding is that cells can maintain polarity without a complete coverage of ECM scaffold around the lobular/tubular structures (Figs. 3 & 5d), suggesting that some cells can be polarized without direct cell-ECM interactions. Nevertheless, without ECM in the medium, cells were unable to form polarity (Fig. 1 and Movie 1). Time-sequence images from confocal microscopy indicate that cells are polarized in small cluster at early stage, and continue expanding into fine 3-D lobular structure under maintained polarization (Fig. 4), which reveals one mechanism of polarization on the entire lobules without complete ECM coverage. Previous studies reported that cadherins from cell-cell junctional adhesions are important in maintaining epithelial structures and establishing epithelial cell polarity(Harris and Peifer, 2004; Jiang et al., 2002; Koumarianou et al., 2017). It is a reasonable hypothesis that those cells without ECM coverage at the top of the lobules/tubules might maintain polarity under cell-cell interactions or intercellular signaling communications, which need be confirmed from further study.

Under suspension culture with addition of soluble ECM, cells developed polarized lobules (with BM) and tubules (with COL) (Fig. 5b), indicating cell-ECM interactions in directing the selforganization of epithelium. Interestingly, multiple lumens were formed on the tubes under suspension culture with COL (Fig. 5e), instead of the homogenous polarity through the tube on solid BM gel (Fig. 1b). This suggests that epithelial tubulogenesis is mediated by ECM microenvironment. Consistently from the previous report, perturbing cell mechanics by inhibition of ROCK-Myosin II pathway also resulted in multiple lumens during tubulogenesis, due to impaired cell motility(Kim et al., 2015). The observations of multi-lumen tubes support that the apicobasal polarization is also mediated in mechanical way beside chemical signals.

In summary, our results highlight the criticalness of cell-ECM interactions in the fluidic phase for epithelial polarization and morphogenesis. Most importantly, we show how a physiological environment can spontaneously emerge through cell-soluble ECM interactions. The gradually collective polarization during tubulogenesis may imply the existence of intercellular communications among the cell group. In the fluidic phase, the polarization occurs in the small cluster at early stage and is maintained through the growth into fine 3-D epithelial structures. These findings might add impact of biophysical factors on how epithelial cells develop and coordinate their polarity and positioning at tissue scales *in vivo,* as well as in the engineering of artificial organs for regenerative medicine *in vitro.*

## Materials and Methods

### Cell Culture, reagents, DNA constructs, and lentivirus

Cell culture medium and reagents were purchased from Invitrogen Gibco. Madin Darby canine kidney (MDCK II) cells (from ATCC) were maintained in Advanced Dulbecco’s modified Eagle’s medium (serum reduced medium) supplemented with 3% fetal bovine serum, 2 mM L-glutamine, 20 unit/ml penicillin, 20 μg/ml streptomycin, and 1 mM sodium pyruvate in a humidified 95% air, 5% CO2 incubator at 37°C.

BD Matrigel (basement membrane matrix, growth factor reduced, phenol red-free), and 3-D Culture Matrix™ Rat Collagen I (5 mg/ml) were purchased from BD Biosciences and R&D Systems, respectively. Rabbit anti-laminin and mouse anti-collagen I primary antibodies were purchased from Sigma, and mouse anti-integrin α6 antibody from Abcam. Pacific Blue-conjugated goat anti-rabbit and anti-mouse IgGs were purchased from Invitrogen, and Rhodamine-conjugated goat anti-mouse IgG antibody from Sigma.

The plasmid pcDNA3-gp135-GFP construct was a gift from Dr. Joachim Füllekrug (Max Planck Institute of Molecular Cell Biology and Genetics)(Meder et al., 2005). The plasmid expressing GFP-tagged human β-actin (GFP-β-actin) under endogenous promoter was provided by Dr. Beat A. Imhof (Switzerland)(Ballestrem et al., 1998). Lentivirus encoding mCherry-conjugated histone H2B (H2B-mCherry) was generously provided by Dr. Rusty Lansford and David Huss (Biology, California Institute of Technology)(Sato et al., 2010).

### Development of stable fluorescent MDCK cell lines

We first developed the stable MDCK cell line expressing H2B-mCherry. Cells at 20-30% confluency were infected with lentivirus encoding H2B-mCherry, and then diluted in 96-well plates to enable the selection and the expansion of single fluorescent colony. Next, we transfected MDCK_H2B-mCherry cells with pcDNA3-gp135-EGFP to develop the cell line expressing both H2B-mCherry and gp135-EGFP (MDCK_H2B-mCherry/gp135-EGFP). In addition, we have transfected MDCK cells with EGFP-β-actin plasmid (MDCK_EGFP-actin). This cell line was used to compare the results obtained from MDCK_H2B-mCherry/gp135-EGFP cells. The transfection was performed by using Lipofectamine2000 (Invitrogen), followed by the antibiotic selection with G418 (300 ug/ml) for 2 weeks to obtain a cell pool displaying various levels of EGFP. A single colony with an intermediate fluorescence intensity of gp135-GFP or GFP-β-actin was selected through dilutions of the cell pool in 96-well plates and further amplified.

### Chambers for cell culture and microscopy

Cell culture experiments and time-lapse microscopy were performed in custom, stainless steel chambers. These chambers were manufactured to have a rectangular shape with a height of 0.6 cm and a 2 × 5.5 cm^2^ internal opening. Nail polish was used to seal 24×60 mm No. 1 cover slips on the bottom of the chambers. To perform multiple-well experiments in one chamber, polydimethylsiloxane (PDMS) blocks were cut to fit the chamber containing multiple wells (~0.5×0.5 cm^2^). The surface of PDMS block was then cleaned and air-dried to allow for a firm attachment on the cover slip of the chamber.

### Cell culture on BM gels or less-adhesive substrates

To make BM gels, we prepared sterile chambers sealed with cover slips on the bottom, and the stock solution (100%) of BD Matrigel was then spread on the top of the cover slips (40-80 μl/cm^2^) followed by incubation at 37°C for 20-30 min. This allowed forming a layer of gel with a variable height (200-400 μm).

To place cells, individual MDCK cells (~1-2×10^4^ cells/cm^2^) were seeded on BM gels in the culture medium containing 2% BM or 20 μg/ml COL. Then, the chamber was placed into petri dish in the cell culture incubator with medium change every 4 days, or every day (or every other day) during time-lapse microscopy. Here, the concentration of BM in the medium followed the “on-top” assay developed by Bissell and her coworkers(Lee et al., 2007). 20 μg/ml COL in the medium was chosen to match the mass concentration of 2% BM (10 mg/ml in stock). Nevertheless, we have applied the different concentrations of COL from 5 to 50 μg/ml, and cells could form tubular structures within this range.

For cell culture on less-adhesive substrates, we first prepared a layer of agarose gel (1%) on the cover slips in the chambers. A mixture of MDCK cells (~10^4^ cells/ml) and culture medium containing 2% BM or 20 μg/ml COL was added into the chambers, and moved in the cell culture incubator. To minimize water evaporation, the chambers were covered by cover slips with a small opening (~5 mm) to allow for air exchange. Medium was changed carefully after 7 days to avoid destroying or losing the cell aggregates.

### Immuno-staining experiments

Immuno-staining experiments were conducted at room temperature, except for incubation with primary antibodies at 4°C. In brief, cell samples were fixed with 4% paraformaldehyde for 15 min, and permeabilized with 0.1% Triton X-100 for 20 min. The cells were then incubated with rabbit anti-laminin, or mouse anti-collagen I at 4°C overnight, followed by incubation with goat secondary antibody conjugated with Pacific Blue (410/455 nm) or Rhodamine (550/570 nm) for 2 hours at room temperature. The images were collected by using epifluorescence or scanning microscopy.

### Epifluorescence and scanning microscopy

Olympus IX71 was equipped with automatic XYZ stage (MS-2000, ASI) and piezo-electric objective stage (P-721 Pifoc, Physik Instrumente) for fast multi-position, z-scanning, and autofocusing time-lapse imaging. An environmental chamber (Haison) was used to maintain humidity, CO2 concentration (5%), and temperature (37°C). For phase-contrast and/or epi-fluorescence microscopy, the imaging system based on Olympus IX71 microscope was equipped with motorized excitation and emission filters with a shutter control (lambda 10-3, Sutter), an Electron-Multiplying CCD camera (ImagEM, C9100-13, Hamamatsu, 512 × 512 pixels, cooled down to −95°C by water circulation), and a 120W fluorescent illumination lamp (X-CITE 120Q, EXFO, Lumen Dynamics Group Inc.). For confocal scanning microscopy, the Olympus IX71 microscope was equipped with lasers of three wavelengths (405nm, 475nm and 594nm), photomultiplier tubes (H10425 and H7422-40, Hamamatsu). The two-photon scanning microscope set up on Olympus IX71 was equipped with a Mai-Tai™ femtosecond laser source (Spectra-Physics). 20x objective (NA: 0.45, Olympus) and 10x objective (NA: 0.3, Olympus) were used in the experiments.

### Image acquisition and analysis

For phase-contrast and/or epifluorescence time-lapse microscopy, Metamorph (version 7.7.3) was used to control the devices and the image acquisition. To acquire z-stack, epifluorescent images were taken at 21 planes with a step size of 4.5 μm. 3-D projection was obtained by collecting pixels with the maximal intensity through the entire z-stack into a single plane by using Metamorph or Matlab programs. For scanning microscopy, Labview was used to run automatic scanning and image acquisition. The exposure time and gains were adjusted according to the image quality and the photo-bleaching effect. Metamorph, Labview, and ImageJ were used for image analysis.

## Acknowledgement

We want to thank Dr. Joachim Füllekrug (Max Planck Institute of Molecular Cell Biology and Genetics) for pcDNA3-gp135-GFP construct, and Dr. Rusty Lansford and David Huss (Biology, California Institute of Technology) for Lentivirus encoding H2B-mCherry. This work is financially supported by Ellison Medical Foundation and Western Heaven Funds (C.G.); National Natural Science Foundation of China (NSFC11872129) and Natural Science Foundation of Jiangsu Province (BK20181416) (M.O.); National Natural Science Foundation of China (NSFC11532003) (L.D.).

## Author Contributions

M. Ouyang and C. Guo designed the research and performed data analysis; M. Ouyang, J-Y. Yu, C. Guo conducted the experiments; J-Y. Yu, Y. Chen, and C. Guo constructed the microscopes and developed the imaging programs; L. Deng provided discussion and certain fund support; M. Ouyang, L. Deng and C. Guo prepared the manuscript.

## Conflict of Interest

the authors declare no conflict of interest in this work.

## Notes

### Competing Interest Statement

The authors have declared no competing interest.

### Summary of Updates

Two major revisions in the updated manuscript: 1) Figures 2 and 3 have been updated in the revised version; 2) there are more additions of contents in the Discussion section.

