## Supporting figure and movies for "Cell-extracellular matrix interactions in the fluidic phase direct the topology and polarity of self-organized epithelial structures"

Supporting information includes:

One Supplementary Figure;

Five Supplementary Movies.

**Running title:** ECM guides the epithelium topology and polarity

### Supplementary Figures

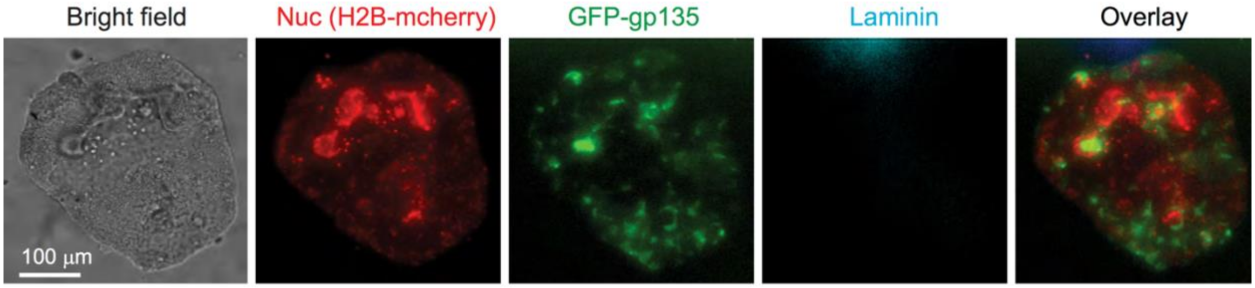


**Figure S1. No lateral assembly of laminin around the non-polarized cell clusters**. MDCK cells were cultured for 12 days on BM gels without soluble BM in the medium, followed by the immuno-staining of laminin. The images show the represented fluorescent images of gp135-GFP (green), H2B-mCherry (red), laminin staining (cyan), and their overlay. Scale bar: 100 μm.

### Supplementary Movie legends

**Movie 1.** Time-lapse epi-fluorescence images of polarized tubule development on a BM gel. MDCK cells expressed H2B-mCherry and gp135-GFP to indicate their nucleus and polarity, respectively. Cells were seeded on a BM gel with 20 μg/ml type I collagen (COL) in the medium. Time interval: 22 minutes.

**Movie 2.** Time-lapse epi-fluorescence images of polarized lobule development on a BM gel. MDCK cells expressed H2B-mCherry and gp135-GFP to indicate their nucleus and polarity, respectively. Cells were seeded on a BM gel with 2% BM in the medium. Time interval: 22 minutes.

**Movie 3.** Time-lapse epi-fluorescence images of random cyst formation on a basement membrane (BM) gel. MDCK cells expressed H2B-mCherry and gp135-GFP to indicate their nucleus and polarity, respectively. Cells were seeded on a BM gel without ECM in the medium. Time interval: 22 minutes.

**Movie 4.** The reconstructed 3-D views of two lobules. Cell nucleus is shown in red color, and laminin immunostaining in green. Note the lateral condensation of laminin around the lobules (outlined by cell nuclei) and the absence of laminin staining on the top and bottom of lobules.

**Movie 5.** Time sequence for the growth of dividing cells into polarized lobule. The MCDK cells expressing gp135-GFP and H2B-mCherry were seeded on BM Matrigel gel with 2% BM in medium, and time-lapse two-photon confocal imaging started with 3 or 4 h interval time one day later (details seen in Figure 4 legend). Acquired confocal images were processed by 3-D reconstructions, and rotated 3-D views of each 4 or 5 time-points (in hours) from the time-sequence images displayed the growth and polarization of the clusters into 3-D lobular structure. As a note, the cell sample on the first two panels may not be exactly the same one on the last four panels, as cells were much motile at early stage during imaging.
